# A putative design for electromagnetic activation of split proteins for molecular and cellular manipulation

**DOI:** 10.1101/2022.11.30.518522

**Authors:** Connor J. Grady, Jory Schossau, Ryan C. Ashbaugh, Galit Pelled, Assaf A. Gilad

## Abstract

The ability to manipulate cellular function using an external stimulus is a powerful strategy for studying complex biological phenomena. One approach to modulate the function of the cellular environment is split proteins. In this method, a biologically active protein or an enzyme is fragmented so that it reassembles only upon a specific stimulus. While there are many tools available to induce these systems, nature has provided other mechanisms that can be utilized to expand the split protein toolbox. Here we show a novel method for reconstituting split proteins using magnetic stimulation. We have found that the Electromagnetic Perceptive Gene (EPG) changes conformation due to magnetic fields stimulation. By fusing split fragments of a certain protein to both termini of the EPG, the fragments can be reassembled into a functional protein under magnetic stimulation due to conformational change. We show this effect with three separate split proteins; NanoLuc, APEX2, and Herpes Simplex Virus Type-1 Thymidine Kinase. Our results show for the first time, that reconstitution of split proteins can be achieved only with magnetic fields. We anticipate that this study will be a starting point for future magnetically inducible split protein designs for cellular perturbation and manipulation. With this technology, we can help to expand the toolbox of the split protein platform and allow better elucidation of complex biological systems.

## Introduction

An ongoing effort in chemical biology is directed towards creating tools that can manipulate cellular systems with molecular precision. A common technique researchers use to address the challenge of temporal and spatial activation of proteins is the split protein approach. This method generally utilizes a functional protein that has been fragmented in a way that allows the split parts to reconstitute and regain native function. Split proteins have been widely used to study protein-protein interactions where researchers fuse split reporter proteins to two interacting proteins and observe their fluorescence^1,2^ or bioluminescence^3,4^. More recently the split protein method has been expanded to control protein function under specific stimulus such as light (optogenetics)^5,6^ or chemicals (chemogenetics)^7^. These approaches have been used to regulate cellular functions such as transcription^8,9^ and enzymatic activity^10,11^ as well as being used in the creation of cellular circuits^12,13^ demonstrating their usefulness for studying complex biological systems.

The current methodologies for controlling split proteins have been well established, nevertheless expanding the split protein toolbox to include tools that respond to different stimuli could lead to further discoveries. One such stimuli with potential for broad impact is magnetism. Magnetic fields represent a noninvasive stimulus with equal distribution that have no limitations of penetration depth. Their “on”/“off” functionality associated with electromagnetic coils also allows for precise control. Several approaches have shown effective magnetic activation of cellular functions by ion channel interactions^14–17^. A novel *electromagnetic perceptive gene* (*EPG*) from the glass catfish *Kryptopterus vitreolus*, has also been shown to have a response to changes in magnetic fields^18–21^.

In this study we combine the split protein approach with the EPG protein to create the first magnetogenetic activatable biological hinge. This system is different from the standard split protein systems where two interacting proteins are needed for the reconstitution of the fragments. Both fragments of the EPG split protein system are fused to either end of the EPG protein and conformational changes in the EPG protein bring the fragments together in the presence of a magnetic field. We tested this with the three established split protein systems of NanoLuc^22^, APEX2^7^, and the Herpes Simplex Virus type 1-Thymidine Kinase (HSV1-TK)^23^. Through these split proteins we showed the potential of this first-of-its-kind technology and set the foundation for future magnetically activated split protein systems.

## Results

### EPG conformation changes in response to magnetic stimulation

Bioluminescence resonance energy transfer (BRET) studies are used for determining if conformational changes occur within a protein^24,25^. Using this design, we studied if EPG has a conformational change due to static magnetic field (10 mTesla). EPG was fused to the blue emitting bioluminescent protein NanoLuc and the yellow emitting fluorescent protein mVenus on the N and C terminals respectively and was expressed in HeLa cells. Fig. 1A shows there is a 2.5% signal increase in the group stimulated by magnetic field over the non-stimulated group. We then designed a BRET construct to test if the protein underwent a dimerization event due to magnetic stimulus. This has the EPG fused to NanoLuc on the C terminal followed by an IRES site followed by EPG fused to mVenus (EPG-NanoLuc IRES EPG-mVenus). The group stimulated with the static magnet had a 1.5% increase compared to the control group (Fig. 1B). The low response implies that dimerization of EPG is not the mechanism by which EPG works. Collectively, these findings suggests that magnetic stimulation led to conformational change of the EPG protein.

**Figure 1:**
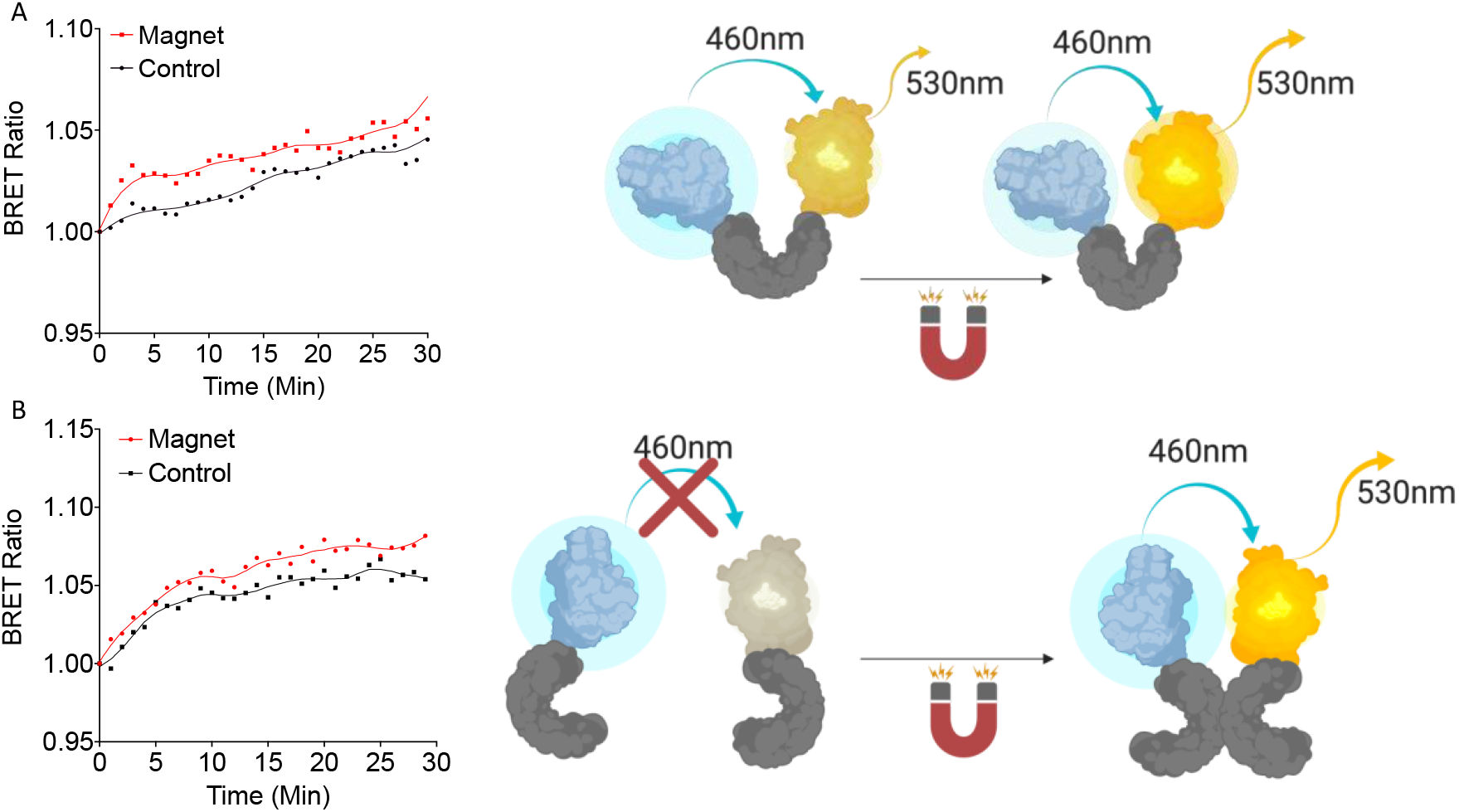
Bioluminescent Resonance Energy Transfer studies of EPG conformational changes. (A) A single copy of EPG cloned between NanoLuc and mVenus. (B) A copy of the EPG was fused to Nanoluc followed by an internal ribosome entry site (IRES) and an EPG fused to an mVenus to express both constructs on the same plasmid. Readings were taken at 530 nm and 460 nm every minute for 30 minutes with or without constant static magnetic stimulation. Readings were normalized to the last read before stimulation. Fit Line in each graph is a Lowess smoothing to show the relationship between the groups. N=15 wells were analyzed for the single and N=9 for the EPG IRES experiments.

### EPG BRET constructs are localized in the cytoplasm

The BRET constructs were cloned in a way that should block the signal sequence and the membrane anchor sequence of the EPG. Therefore, we anticipated cytoplasmic expression. To test this hypothesis, we co-expressed the EPG BRET construct as well as an EPG HaloTag construct that was previously shown to be membrane anchored in mammalian cells^26^. Fluorescent images show the BRET construct was likely expressed in the cytoplasm as opposed to the EPG HaloTag fusion protein that is mostly observed on the cellular membrane. Fig. 2 demonstrates the EPG BRET construct to be a cytoplasmic protein providing evidence to support that the membrane and signal sequences were blocked. The conformational change that occurs in the cytoplasm also indicates that the magnetoreception of EPG is not dependent of its cellular localization.

**Figure 2:**
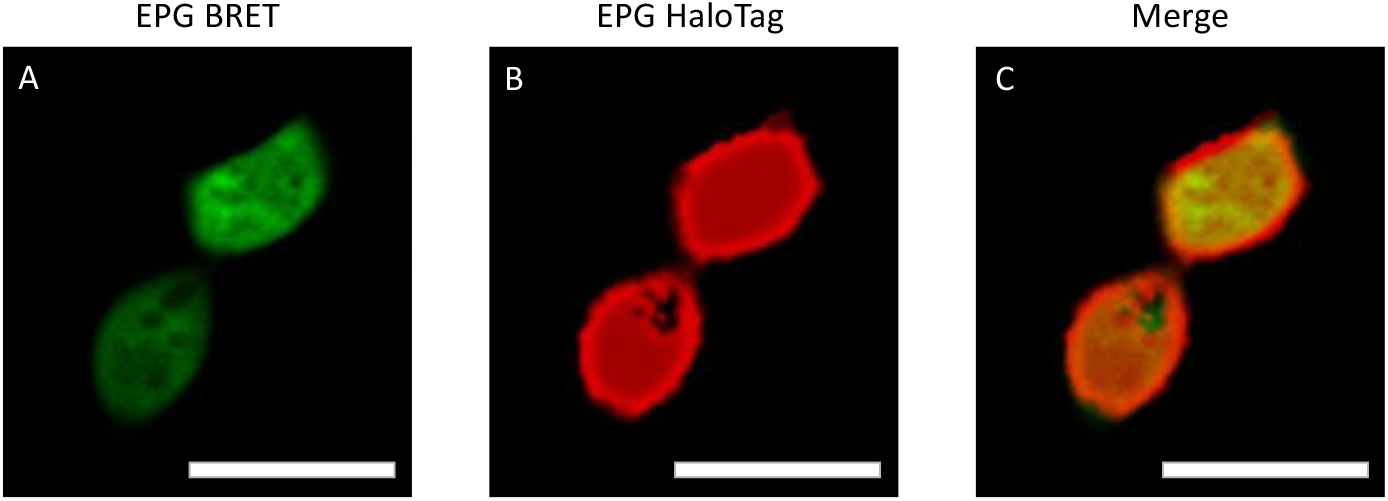
EPG BRET Fluorescent Imaging for Cell Localization. (A) Hela cells cotransfected with EPG BRET construct imaged with the GFP filter and (B) EPG HaloTag labeled with Janelia Fluor 646 dye imaged with Cy5 filter. (C) A merged image of the two channels shows expression of the EPG BRET construct in the cytoplasm and the EPG HaloTag construct on the cell membrane. Scale bar = 25 μm.

### EPG split NanoLuc expressed in E. Coli exhibits an increase in bioluminescence in response to magnetic stimulation

Building upon the split protein concept and on the magnetoresponsive properties of the EPG, we have developed a new platform that allows remote activation of a protein or enzyme using electromagnetic fields (EMF). The principle for this tool is cloning the EPG between two parts of a split protein or between two enzymes/proteins that need close proximity for activation (Fig. 3A). Here we split NanoLuc (171 amino acids) into two fragments at amino acid sites 65 and 66. The 1-65 and 66-171 fragments were fused to the N and C termini of EPG respectively. We chose this split site based on previous reports^22^ (Fig. 3B). A truncated version of this construct was created by removing the signal sequence and membrane anchor sequence of EPG. Another construct was created by using the reverse nucleotide sequence of the truncated EPG and this was referred to as flipped trEPG. When exposed to EMF, the EPG construct in the intact cells showed up to 68.7±24.6% increase in luminescence in contrast to controls constructs (Fig. 3C). Under the same condition but when measured in cell extract, the EPG construct displayed a 39.4±41.4% compared to control truncated or reverse truncated EPG (Fig. 3D). These results are the first demonstration that a split protein can be brought together by the conformational change of EPG. Thus, EPG can act as a magnetically activatable hinge.

**Fig. 3:**
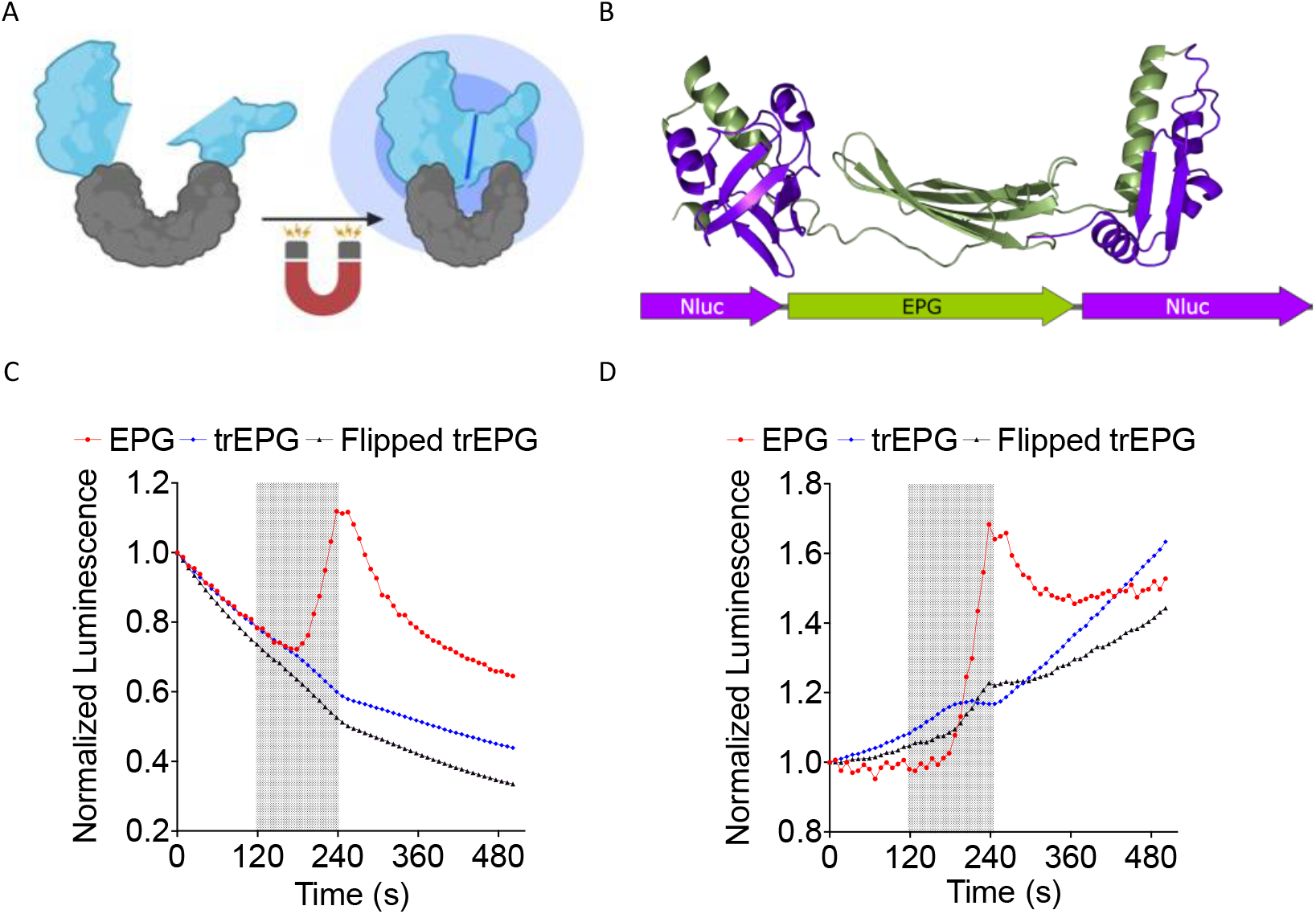
EPG split NanoLuc experiments in E. coli BL21 cells. Readings were taken on the IVIS every 10 seconds with an open filter. Electromagnetic stimulus was applied to the cells for 2 minutes and shown as shaded region. (A) Illustration of EPG split NanoLuc construct. (B) A model of the EPG split NanoLuc construct. E. coli Lysate (C) and whole cell E. coli (D) containing EPG split NanoLuc showed an increase in luminescence in contrast to EPG truncated and Flipped EPG. Results shown are duplicate experiments with N=6 in each trial.

### Remote magnetic control of peroxidase activity using EPG split APEX2

To demonstrate that the EPG split approach can be used as a platform technology, we used a Split APEX2 Peroxidase^7^. This system allows simplified demonstration of the concept that EMF can control an enzymatic reaction and the output can be measured directly with colorimetric or fluorescent reaction with any standard plate reader or potentially even a microscope. HEK 293FT cells expressing EPG split APEX2 treated with both static magnetic stimulus and hydrogen peroxide displayed a clear increase in fluorescence (150±16%; Fig. 4) compared to the cells that did not experience magnetic stimulation. These results show a statistically significant increase in peroxidase activity in response to 30 minutes of exposure to static magnetic field. We also repeated this experiment at room temperature and 37°C and found similar results (Fig. S1). These findings indicate that the EPG protein can be used as magneto-switch to activate multiple enzymes.

**Fig. 4:**
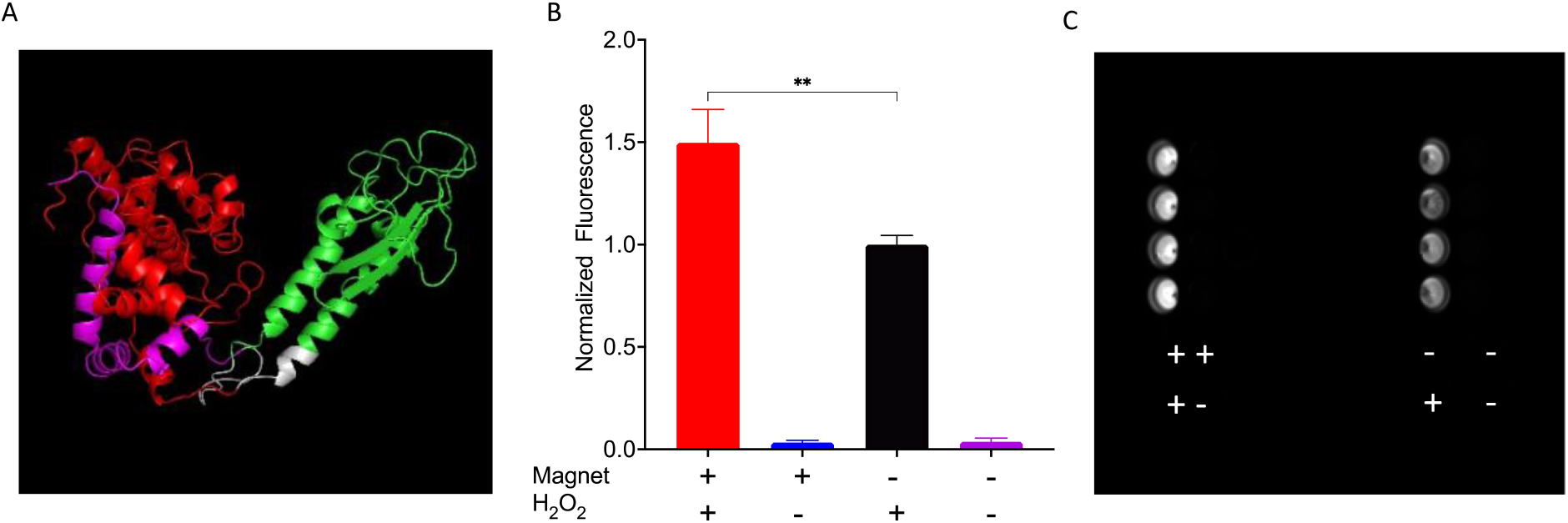
HEK 293FT cells expressing EPG split APEX2 show an increase in fluorescence in response to magnetic field. All wells were treated with Amplex UltraRed reagent and the four combinations of with or without magnetic stimulus and H2O2 for 30 minutes. (A) Predicted structure of EPG split APEX2 with EPG (green), AP fragment (Red), EX fragment (magenta), and linkers (white). (B) Endpoint results of cells treated with all combinations of static magnetic stimulus and hydrogen peroxide (n=4 independent experiments). (C) Image of a plate taken with Cy3 filter after experiment for detection of resorufin accumulation.

### EPG split HSV1-TK Ganciclovir-Mediated Cell Death

To demonstrate potential therapeutic and diagnostic usage of an EPG split protein, we decided to create an EPG split herpes simplex virus thymidine kinase. This construct was based on the previously split sr39 mutant of HSV1-TK^23^. To test the EPG split HSV-TK we used the proven suicide gene therapy combination of HSV1-TK and ganciclovir (GCV). Where the HSV-TK phosphorylates the GCV allowing other cellular enzymes to further phosphorylate and incorporate the GCV triphosphate into the DNA causing cell death^27^. To perform the cell uptake assay of the GCV, 4T1-Luc2 cells transfected with either EPG split HSV1-TK, wildtype HSV1-TK or a mock transfection were subjected to 0.15 mg/ml of GCV for 72 hours. The 4T1-Luc2 cell line expresses the gene Luc2, an ATP dependent luciferase. Therefore, cell viability was quantified by directly measuring luminescence after the 72 hours of incubation with GCV (Fig. 5A). A linker screening was performed with three combinations of an EPG without the signal sequence and membrane localization signal and four full length EPG constructs. These constructs were cloned with either flexible (GGGGS) or rigid (PAPAP) linkers between the EPG and split fragments of the HSV1-TK and then were tested for magnetic response after treatment with GCV. After the initial screening two constructs showed potential and the full-length EPG with a flexible first linker and rigid second linker was chosen due to a lower basal activity (Fig. S2).

**Figure 5:**
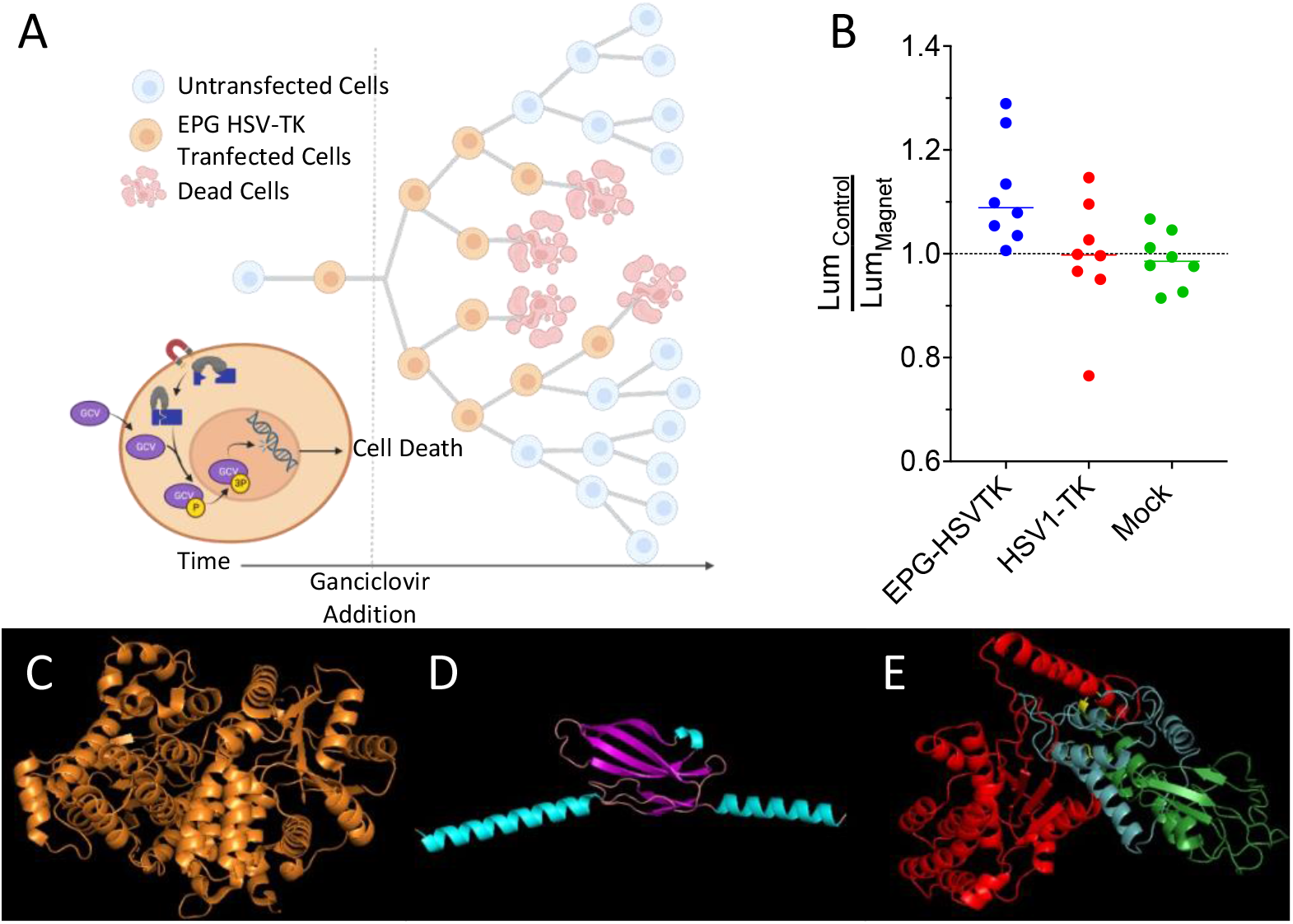
Ganciclovir Mediated Cell Death; Control vs Magnet. (A) Schematic of the experimental process and design. (B) The ratio of average control cell luminescence to magnetic stimulated cell luminescence over the course of eight experimental replicates. (C) Structure of HSV1-TK; (D) predicted structure of EPG with core structure (purple) and signal sequence and membrane anchor sequence (teal), (E) and predicted structure of EPG split HSV1-TK with N-terminal HSV1-TK (red), EPG (green), and C-terminal HSV1-TK (blue).

To test this construct further, eight experimental replicates were performed comparing magnetic stimulated cells and non-stimulated cells. In each of these replicates the mean of luminescence of the EPG-HSV1-TK magnetic stimulated cells was lower than the control EPG-HSV1-TK cells with the average percent change between these groups of 10% (Figs. 5B and S3 A-H), the probability for such event is 0.00039; Supplementary material and Figs. SI4). This was not the case with the HSV1-TK (Figs. 5B and S3 I-P) and mock transfected (Figs. 5B and S3 Q-X) groups which have an average of 3.6% and 1.3% respectively. In both these control groups there was no consistent trend of cell viability due to magnetic stimulation as both groups showed three experiments with lower average cell viability and five experiments of increased in cell viability in the presence of magnetic field (Fig 5B; the probability for such event is 0.375; Supplementary material and Figs. SI4). Therefore, it appears that even in a complex system such as EPG split HSV1-TK and GCV, a significant yet small effect of magnetic field can be measured. Together with the other experiments, our finding implies that EPG can be used as a bio-magnetic switch for remote magnetic activation of enzymes.

## Discussion

We have demonstrated a novel method to control split proteins using magnetic fields. To our knowledge, we are the first to create this kind of technology. We were able to see an effect of the magnetic field in each of the constructs shown, but challenges arose in some of the constructs. For example, in the EPG HSV1-TK experiments, the high number cell divisions in each of the experimental replicates led to small measurable effect. Because of the design of this experiment many factors attributed to this high number of cell divisions^27^ (Fig. 5A). Since this is a suicide gene therapy, the number of starting cells highly influenced the outcome of this experiment. This coupled with transient transfection and the fact that ganciclovir incorporates during cell division causes non-transfected cells to take over the cell population. Despite this, we still see a significant effect of the magnetic stimulated groups compared to the control groups. Nevertheless, using remote controlled HSV1-TK has a great therapeutic and diagnostic importance for positron emission tomography (PET)^28,29^ and magnetic resonance imaging (MRI)^30,31^. Further optimization of the EPG, the linker using HSV1-TK mutants^32^ or using alternative nucleoside kinases^33,34^ might result in improved performance of the construct. This will potentially allow imaging in in vivo systems.

Interestingly, we found that in the split NanoLuc experiments, keeping the signal sequence and membrane anchor sequence is critical. Even though those sequences do not play a role in the magnetoreception, as they are believed to be removed in the post translational processing of EPG, they act as linker sequences for the split proteins. When removed, the split protein did not respond to magnetic fields and provided a much higher luminescence. Thus, we theorize that the split halves have reassembled independent of the function of the EPG. Similar phenomena were previously reported for some split proteins^35^. Because of the hinge design of these constructs, the close proximity of split halves needs to be considered during the design process.

This work creates a new platform to control split proteins. This study was focused on establishing the platform rather than optimizing each individual construct created. The constructs other than EPG HSV1-TK, went through no linker screening and were mainly based on existing linkers used with the split proteins utilized. While this is one aspect that should be optimized for future studies, we were still able to see the magnetic effect in each of these constructs. Therefore, this work sets the future of using magnetic fields as an induction mechanism to control split proteins.

## Materials and Methods

### Plasmid Construction

See Supplementary material for a list of constructs used in this study. For cloning, PCR fragments were amplified using Platinum Superfi II Polymerase (Invitrogen). The assembly of fragments was done by either NEBuilder HiFi Assembly (New England BioLabs, NEB) or TOPO directional cloning (Invitrogen). Assembled products were heat-shock transformed to 5-alpha (NEB), TOP10 (Invitrogen) or BL21 Star™ (DE3) (Invitrogen) bacteria.

### Bioluminescent Resonance Energy Transfer Assay

HeLa cells were split to 70% confluency in a 6 well plate. The following day cells were transfected with plasmid DNA according to Lipofectamine 3000 protocol. The transfection efficiency was checked under the Keyence microscope using the GFP filter. Cells were then split to black walled clear bottom plastic 96 well plates. A stock solution (50 mM) of h-Coelenterazine (h-CTZ, NanoLight Technologies) was prepared by adding 25uL of solution to dried h-CTZ powder. A working concentration of 5uM was made by diluting the h-CTZ stock solution in FluoroBrite DMEM (Gibco).

Prior to measurements, culture media was aspirated from cells and replaced with h-CTZ containing media. The plate was then put into a Victor Nivo (Perkin Elmer) plate reader. Reads were taken every minute for 15 minutes from the bottom of the plate using 480/30nm and 540/30nm filters. The plate was then taken out and static magnets were put into wells for magnet samples and then the plate was placed back in the reader and readings were taken every minute for 30 minutes. A ratio of the 540/480 was used to calculate BRET efficiency.

### NanoLuciferase Assay

Plasmids containing NanoLuciferase constructs were transformed into BL21 *E. coli* cells. Colonies were picked and grown in Magic Media (Invitrogen) expression media overnight at 37°C. After overnight expression, cells were pelleted by centrifugation followed by resuspension in PBST and were sonicated using 10 sec on 20 second on pulses for 2-3 minutes to create cell lysates.

For IVIS (Perkin Elmer) imaging 25 uL of cells or cell lysate were added to the 96 well plate followed by 150uL of LB broth with 5uM h-CTZ. 15 min after the addition of h-CTZ, IVIS images were captured using a 1 second exposure time with an open emission filter and an F stop of 1 which allowed us to capture an image every 10 seconds. After 2 minutes of imaging, an electromagnetic coil (35 mTesla field strength) surrounding the 96 well plate^36^ was turned on and samples were under electromagnetic stimulation for a 2 minute period at which the magnet was turned off and images were captured for another 6 minutes. Images were analyzed using the Living Image Software (Perkin Elmer).

### Amplex Ultrared Assay

HEK 293FT cells were grown to 70-90% confluency and transfected in a 6 well plate according to manufacturer’s protocol (Lipofectamine 3000). After 24 hours post transfection, cells were split into black walled 96 well plates and left for to grow for 18-24 hours. Cells were then moved to ice and media was replaced with a solution of 50uM Amplex UltraRed (Life Technologies) with 0.02% (6.7mM) H202 in PBS. Cells with magnet stimulation had static magnets (150-200 mTesla) on top and bottom of well plate over the stimulated wells. Stimulation occurred for 30 minutes and then were read on Cytation 5 plate reader (BioTek) using 530 excitation and 590 emission read settings. For image (Fig 4B) taken on Chemidoc (BioRad) using the Cy3 blot function.

### Ganciclovir Mediated Cell Death

4T1 Luc2 (ATCC) cells were plated at 10,000-20,000 cells per well into 96 well plates. After 8 hours, cells were transfected according to manufacturer’s protocol (Lipofectamine 3000). The following day media was exchanged with media containing 0.15mg/mL ganciclovir (InvivoGen). Magnet stimulated cells were then placed under constant magnetic stimulation for 72 hours. After 72 hours viability was measured by exchanging media with Fluorobrite (Invitrogen) supplemented with 0.15 mg/mL d-Luciferin (Gold Biotechnology). Luminescent reads were then taken on a Spark (Tecan) plate reader.

### Software

Creation of the protein models were performed using the Robetta server^37^. All models were predicted using the RoseTTAFold modeling method. Illustrations used in figures were created with BioRender.com. Graphs and statical analysis were performed using GraphPad Prism version 9.4.1 for macOS, GraphPad Software, San Diego, California USA, www.graphpad.com.

## Acknowledgements

The authors would like to thank Brianna Ricker for critical editing of the manuscript. A.A.G acknowledges financial support from the NIH/NINDS: R01-NS098231; R01-NS104306 NIH/NIBIB: R01-EB031008; R01-EB030565; R01-EB031936 and NSF 2027113.

## References

1. Romei, M.G. & Boxer, S.G. Split Green Fluorescent Proteins: Scope, Limitations, and Outlook. Annu Rev Biophys 48, 19–44 (2019).

2. Hu, C.D. & Kerppola, T.K. Simultaneous visualization of multiple protein interactions in living cells using multicolor fluorescence complementation analysis. Nat Biotechnol 21, 539–45 (2003).

3. Paulmurugan, R. & Gambhir, S.S. Monitoring protein-protein interactions using split synthetic renilla luciferase protein-fragment-assisted complementation. Anal Chem 75, 1584–9 (2003).

4. Dixon, A.S. et al. NanoLuc Complementation Reporter Optimized for Accurate Measurement of Protein Interactions in Cells. ACS Chem Biol 11, 400–8 (2016).

5. Yu, Y. et al. Engineering a far-red light-activated split-Cas9 system for remote-controlled genome editing of internal organs and tumors. Sci Adv 6, eabb1777 (2020).

6. Kawano, F., Okazaki, R., Yazawa, M. & Sato, M. A photoactivatable Cre-loxP recombination system for optogenetic genome engineering. Nat Chem Biol 12, 1059–1064 (2016).

7. Han, Y. et al. Directed Evolution of Split APEX2 Peroxidase. ACS Chem Biol 14, 619–635 (2019).

8. Zetsche, B., Volz, S.E. & Zhang, F. A split-Cas9 architecture for inducible genome editing and transcription modulation. Nat Biotechnol 33, 139–42 (2015).

9. Nihongaki, Y., Otabe, T., Ueda, Y. & Sato, M. A split CRISPR-Cpf1 platform for inducible genome editing and gene activation. Nat Chem Biol 15, 882–888 (2019).

10. Cubillos-Ruiz, A. et al. An engineered live biotherapeutic for the prevention of antibiotic-induced dysbiosis. Nat Biomed Eng 6, 910–921 (2022).

11. Shekhawat, S.S. & Ghosh, I. Split-protein systems: beyond binary protein-protein interactions. Curr Opin Chem Biol 15, 789–97 (2011).

12. Gao, X.J., Chong, L.S., Kim, M.S. & Elowitz, M.B. Programmable protein circuits in living cells. Science 361, 1252–1258 (2018).

13. Fink, T. et al. Design of fast proteolysis-based signaling and logic circuits in mammalian cells.

14. Huang, H., Delikanli, S., Zeng, H., Ferkey, D.M. & Pralle, A. Remote control of ion channels and neurons through magnetic-field heating of nanoparticles. Nat Nanotechnol 5, 602–6 (2010).

15. Stanley, S.A., Sauer, J., Kane, R.S., Dordick, J.S. & Friedman, J.M. Remote regulation of glucose homeostasis in mice using genetically encoded nanoparticles. Nat Med 21, 92–98 (2015).

16. Stanley, S.A. et al. Radio-wave heating of iron oxide nanoparticles can regulate plasma glucose in mice. Science 336, 604–8 (2012).

17. Wheeler, M.A. et al. Genetically targeted magnetic control of the nervous system. Nat Neurosci 19, 756–761 (2016).

18. Krishnan, V. et al. Wireless control of cellular function by activation of a novel protein responsive to electromagnetic fields. Sci Rep 8, 8764 (2018).

19. Metto, A.C., Gilad, A.A. & Pelled, G. Magnetogenetic closed-loop reduction of seizure activity in a rat model of epilepsy. bioRxiv, 2022.08.19.504501 (2022).

20. Cywiak, C. et al. Non-invasive neuromodulation using rTMS and the electromagnetic-perceptive gene (EPG) facilitates plasticity after nerve injury. Brain Stimul 13, 1774–1783 (2020).

21. Hwang, J. et al. Regulation of Electromagnetic Perceptive Gene Using Ferromagnetic Particles for the External Control of Calcium Ion Transport. Biomolecules 10(2020).

22. Zhao, J., Nelson, T.J., Vu, Q., Truong, T. & Stains, C.I. Self-Assembling NanoLuc Luciferase Fragments as Probes for Protein Aggregation in Living Cells. ACS Chem Biol 11, 132–8 (2016).

23. Massoud, T.F., Paulmurugan, R. & Gambhir, S.S. A molecularly engineered split reporter for imaging protein-protein interactions with positron emission tomography. Nat Med 16, 921–6 (2010).

24. Audet, N. et al. Bioluminescence resonance energy transfer assays reveal ligand-specific conformational changes within preformed signaling complexes containing delta-opioid receptors and heterotrimeric G proteins. J Biol Chem 283, 15078–88 (2008).

25. Inagaki, S. et al. Genetically encoded bioluminescent voltage indicator for multi-purpose use in wide range of bioimaging. Sci Rep 7, 42398 (2017).

26. Mitra, S., Barnaba, C., Schmidt, J., Pelled, G. & Gilad, A.A. Functional Characterization of an Electromagnetic Perceptive Protein. bioRxiv, 2020.10.07.329946 (2020).

27. Alon, L. et al. Molecular Imaging of CXCL12 Promoter-driven HSV1-TK Reporter Gene Expression. Biotechnology and Bioprocess Engineering 23, 208–217 (2018).

28. Serganova, I., Ponomarev, V. & Blasberg, R. Human reporter genes: potential use in clinical studies. Nucl Med Biol 34, 791–807 (2007).

29. Keu, K.V. et al. Reporter gene imaging of targeted T cell immunotherapy in recurrent glioma. Sci Transl Med 9(2017).

30. Bar-Shir, A., Liu, G., Greenberg, M.M., Bulte, J.W.M. & Gilad, A.A. Synthesis of a probe for monitoring HSV1-tk reporter gene expression using chemical exchange saturation transfer MRI. Nat. Protocols 8, 2380–2391 (2013).

31. Bar-Shir, A. et al. Transforming thymidine into a magnetic resonance imaging probe for monitoring gene expression. J Am Chem Soc 135, 1617–1624 (2013).

32. Allouche-Arnon, H. et al. Computationally designed dual-color MRI reporters for noninvasive imaging of transgene expression. Nat Biotechnol (2022).

33. Likar, Y. et al. A new pyrimidine-specific reporter gene: a mutated human deoxycytidine kinase suitable for PET during treatment with acycloguanosine-based cytotoxic drugs. J Nucl Med 51, 1395–403 (2010).

34. Bar-Shir, A. et al. Quantification and tracking of genetically engineered dendritic cells for studying immunotherapy. Magnetic Resonance in Medicine 79, 1010–1019 (2018).

35. Dolberg, T.B. et al. Computation-guided optimization of split protein systems. Nat Chem Biol 17, 531–539 (2021).

36. Ashbaugh, R.C., Udpa, L., Israeli, R.R., Gilad, A.A. & Pelled, G. Bioelectromagnetic Platform for Cell, Tissue, and In Vivo Stimulation. Biosensors (Basel) 11(2021).

37. Baek, M. et al. Accurate prediction of protein structures and interactions using a three-track neural network. Science 373, 871–876 (2021).

