## Supplemental Information for "A putative design for electromagnetic activation of split proteins for molecular and cellular manipulation"

**Supplemental Figures**

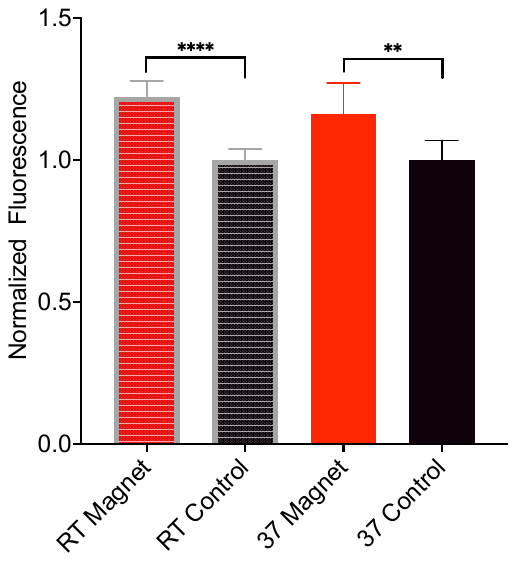

**Supplemental Figure 1: EPG split APEX2 temperature variation**.

Comparison of the EPG split APEX2 system at room temperature and 37C. Cells were either subjected to magnetic field (red) or no stimulus (black). N=8 biological replicates per group. The (**) denotes p-value <0.01 and the (****) denotes a p-value <0.0001.

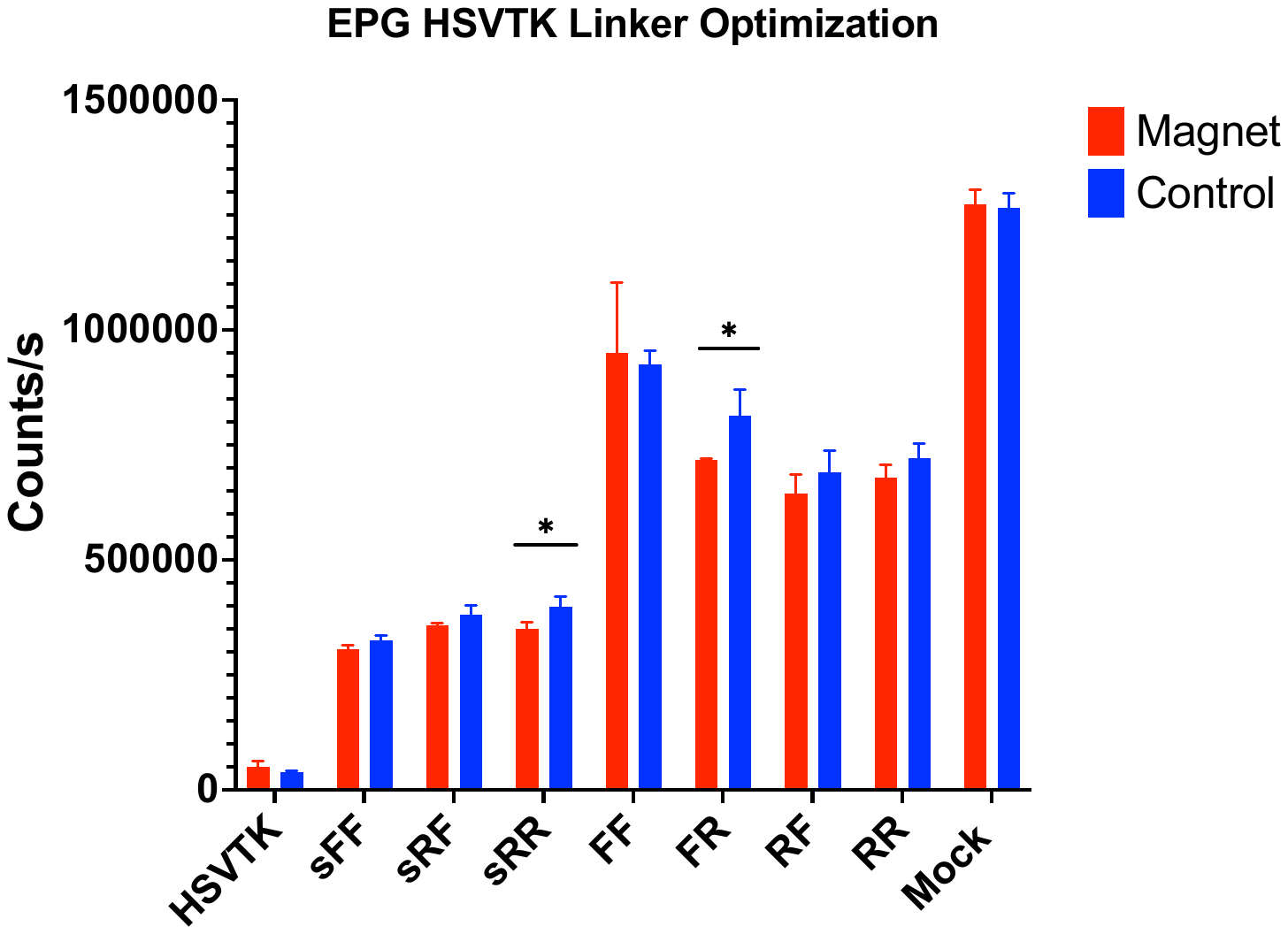
**Supplemental Figure 2: Linker Screening of EPG-HSVTK constructs in 4T1 Cells.**

Bioluminescent readouts from constructs after 72 hours of GCV treatment with and without magnetic stimulation. The lowercase “s” denotes EPG without signal sequence and membrane sequences. The capital “F” denotes a flexible linker and capital “R” denotes a rigid linker. The “*” denotes a p-value <0.05.

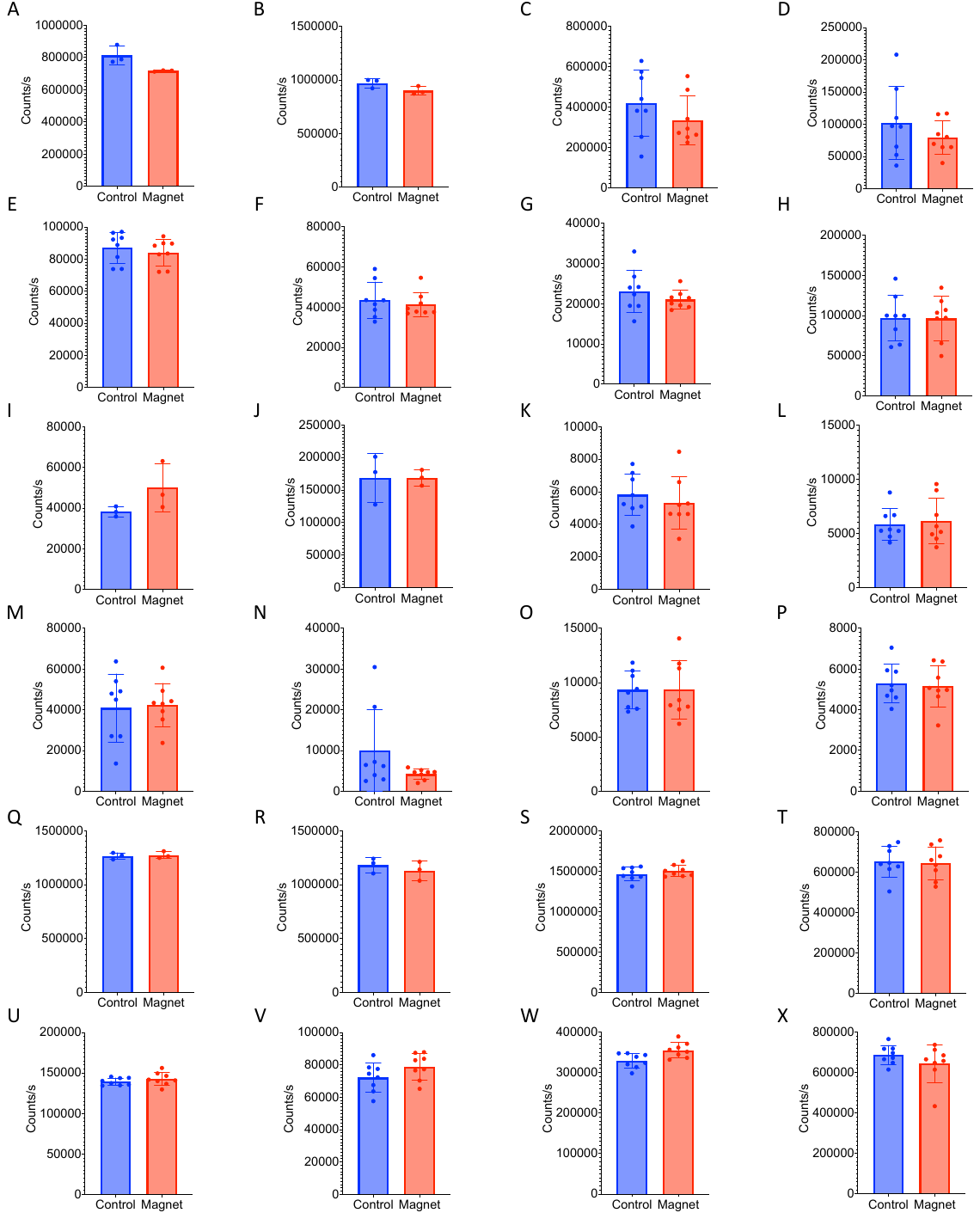

**Supplemental Figure 3: Ganciclovir mediated cell death in 4T1 cells**.

Cells expressed the EPG split HSV1-TK construct (A-H), HSV1-TK construct (I-P), or were mock transfected (Q-X) and cell viability in either a magnetic stimulated or control conditions.

**Statistical analysis of the HSV1-TK experiments:**

The experiment includes 8 replicates from which we observed that the average cell growth was inhibited when under magnetic influence compared to the non-magnetic condition, and compared to each of 8 replicates of controls expecting cell growth, and cell death. There are 2 questions about the statistical significance of this observation: 1) “How likely is it to again observe the 8 replicate outcomes of reduced average cell growth if the magnetic condition actually had no effect?” 2) “How likely is it to again observe in each replicate the particular difference of averages, if the magnetic condition actually had no effect?” These can be thought of as condition and replicate significance testing, respectively.

For the condition significance testing, we investigated how likely it is to observe all 8 replicates of average cell growth showing inhibition if the experimental magnetic condition had no effect. To test this, we simulated how often we observe all 8 replicates with inhibited cell growth if we were to perform the growth and death control replicates many more times. This test is simulated because more replicate data becomes costly and laborious to gather. To simulate more control replicates we sampled our existing control replicates with replacement, counting how many samples of 8 contain all 8 showing inhibited cell growth. By chance, we observe our experimental results from the simulated sampling of the control conditions with a probability of 0.00039 (see Equation 1). This is sufficiently low to suggest significance of inhibition between replicate conditions. Probability density for the binomial distribution is shown in Equation 1, where n is the number of trials, p is the probability of success, and N is the number of successes.

$$P\left( N \right)=\left( \begin{matrix} n \\ N \end{matrix} \right)p^{N}{(1-p)}^{n-N}$$

Equation: 1. Our control conditions both have 3 replicates with average inhibited cell growth or death, and 5 replicates with invigorated cell growth or death. Assuming this outcome was the most common outcome to observe of the underlying distribution, then the probability of inhibition in the controls then becomes 3/8 or p=0.375, and N=8. Using the binomial formula yields P(8)=0.00039.

For the replicate significance testing, we investigated how likely it is to observe, for each replicate, the particular difference of means if the experimental magnetic condition had no effect and the data for both conditions had come from the same distribution. To test this, we used a Randomization Test^1^ wherein the data comprising each replicate for both magnetic and non-magnetic conditions are assumed to originate from the same source. We randomize the “magnetic” and “non-magnetic” labels from the collected data for each replicate, then recalculate the difference of means. When performed many times (1,000,000), this process creates a distribution of differences, from which we can calculate how often the actual observed means difference or better can arise. All experimental condition replicates (EPG-HSV1-TK) showed statistical significance that the observed difference of means was far greater than the 95% confidence interval of the mean in which the differences of the means were sampled from randomized data assuming no effect of the magnetic condition (Figures S4-S6).

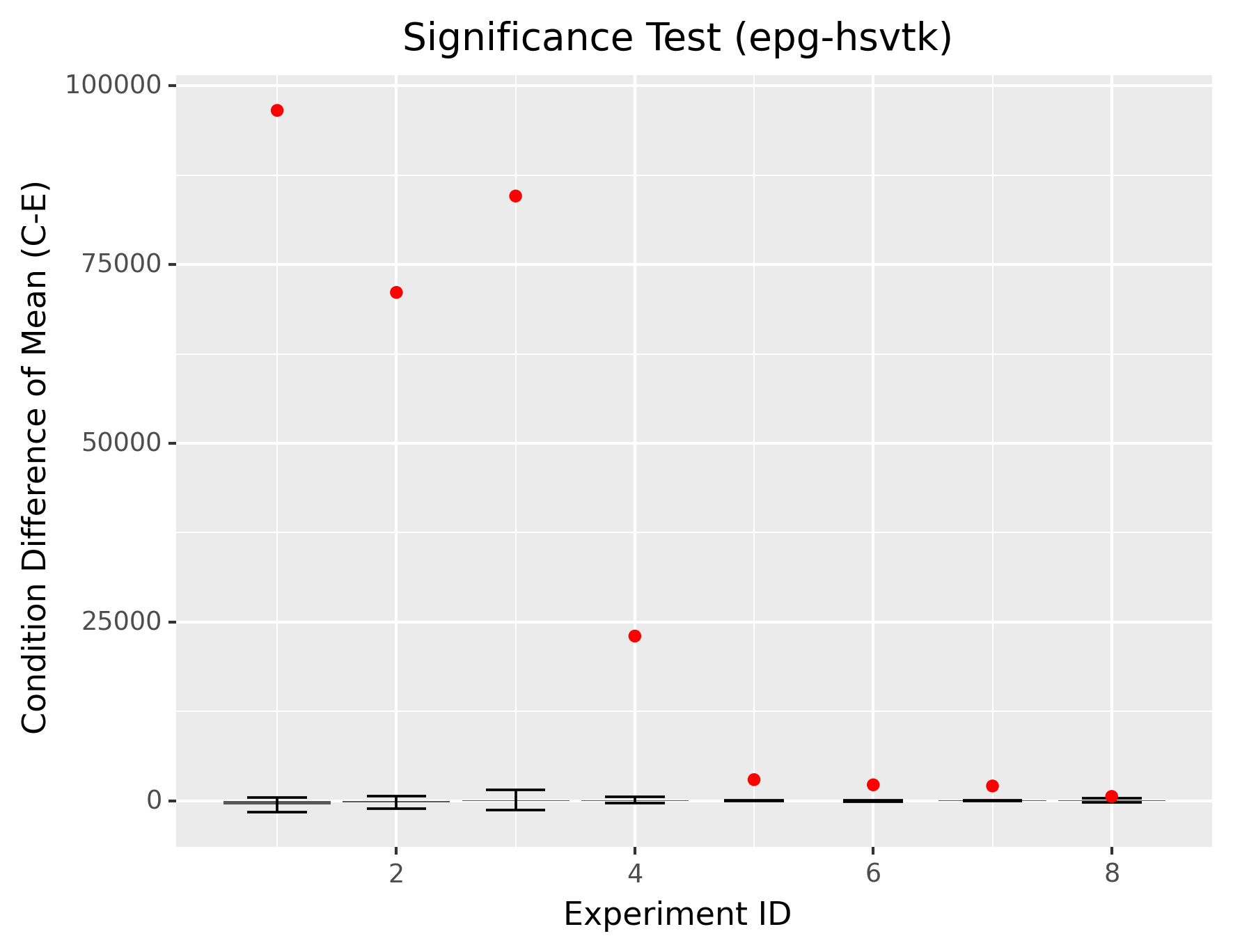
**Supplemental Figure 4: Significance Testing of EPG-HSV1-TK construct for Ganciclovir mediated cell death in 4T1 cells**.

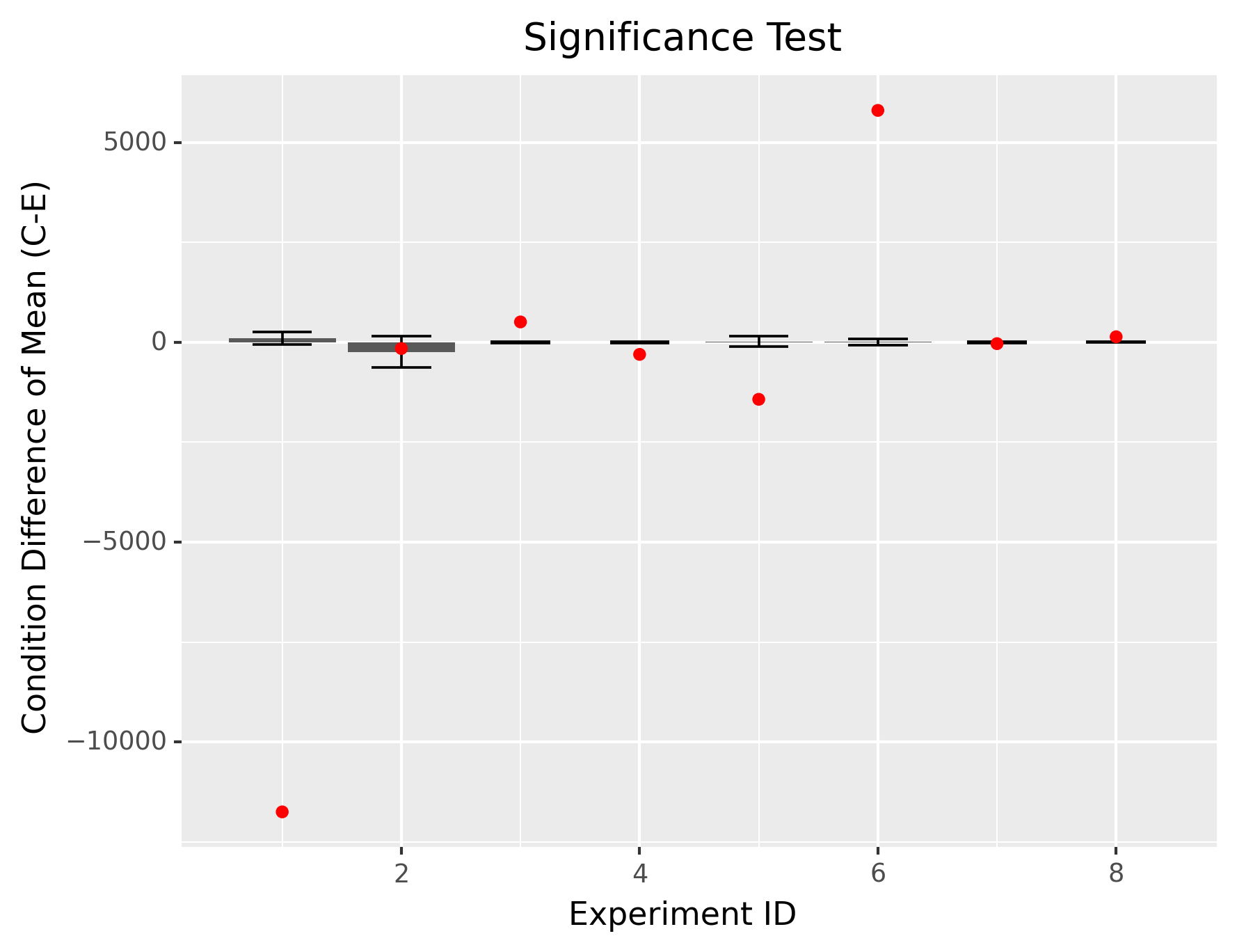

**Supplemental Figure 5: Significance Testing of HSV1-TK construct for Ganciclovir mediated cell death in 4T1 cells.**

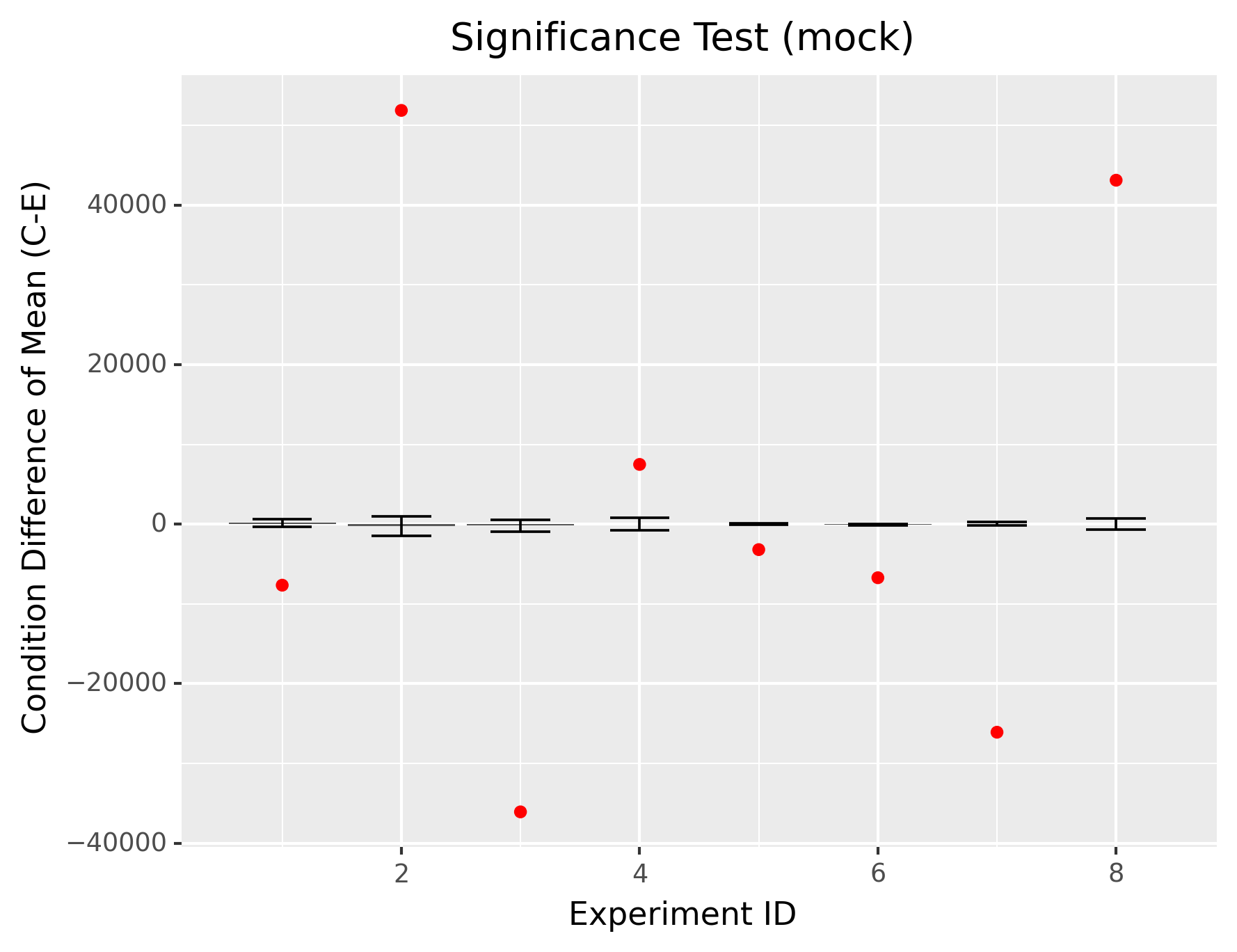

**Supplemental Figure 6: Significance Testing of Mock transfected cells for Ganciclovir mediated cell death in 4T1 cells.**

Constructs

**EPG BRET Constructs**

**EPG BRET- Vector**: pcDNA 3.1 Promoter: CMV

Start-NanoLuc-EPG-mVenus-Stop

ATGGTCTTCACACTCGAAGATTTCGTTGGGGACTGGCGACAGACAGCCGGCTACAACCTGGACCAAGTCCTTGAACAGGGAGGTGTGTCCAGTTTGTTTCAGAATCTCGGGGTGTCCGTAACTCCGATCCAAAGGATTGTCCTGAGCGGTGAAAATGGGCTGAAGATCGACATCCATGTCATCATCCCGTATGAAGGTCTGAGCGGCGACCAAATGGGCCAGATCGAAAAAATTTTTAAGGTGGTGTACCCTGTGGATGATCATCACTTTAAGGTGATCCTGCACTATGGCACACTGGTAATCGACGGGGTTACGCCGAACATGATCGACTATTTCGGACGGCCGTATGAAGGCATCGCCGTGTTCGACGGCAAAAAGATCACTGTAACAGGGACCCTGTGGAACGGCAACAAAATTATCGACGAGCGCCTGATCAACCCCGACGGCTCCCTGCTGTTCCGAGTAACCATCAACGGAGTGACCGGCTGGCGGCTGTGCGAACGCATTCTGGCGATGAAGTGTGTACTTTTGGGATTCGCAGCAGTGATCGGATTCTTCGCGATCGCGGAGTCTCTTACCTGTAACACATGCTCAGTGAGTCTGATTGGAATATGTCTGAATCCCGCAACAGCGACTTGCTCCACCAACACATCCGTCTGCACCACAGGAAGAGCCAGTTTCACGGGCGTCCTCGGCTTCCTGGGCTTCAACTCCCAGGGCTGCACGGAGGGAGCTCAGTGTAATGGCACCGTGTCCGGGTCCATCCTGGGTGCGTCGTACACGGTCACTCAAACCTGCTGCAGCACAAACAACTGCAACCCCGTGACCAGCGGCGCCTCCTACGTCCAGATCTCCGTCAGCGCGGCCCTGAGCGCCGCCCTGCTGGCCTGCGTCTGGGGCCAGTCCGTCTACATGGTGAGCAAGGGCGAGGAGCTGTTCACCGGGGTGGTGCCCATCCTGGTCGAGCTGGACGGCGACGTAAACGGCCACAAGTTCAGCGTGTCCGGCGAGGGCGAGGGCGATGCCACCTACGGCAAGCTGACCCTGAAGCTGATCTGCACCACCGGCAAGCTGCCCGTGCCCTGGCCCACCCTCGTGACCACCCTGGGCTACGGCCTGCAGTGCTTCGCCCGCTACCCCGACCACATGAAGCAGCACGACTTCTTCAAGTCCGCCATGCCCGAAGGCTACGTCCAGGAGCGCACCATCTTCTTCAAGGACGACGGCAACTACAAGACCCGCGCCGAGGTGAAGTTCGAGGGCGACACCCTGGTGAACCGCATCGAGCTGAAGGGCATCGACTTCAAGGAGGACGGCAACATCCTGGGGCACAAGCTGGAGTACAACTACAACAGCCACAACGTCTATATCACCGCCGACAAGCAGAAGAACGGCATCAAGGCCAACTTCAAGATCCGCCACAACATCGAGGACGGCGGCGTGCAGCTCGCCGACCACTACCAGCAGAACACCCCCATCGGCGACGGCCCCGTGCTGCTGCCCGACAACCACTACCTGAGCTACCAGTCCGCCCTGAGCAAAGACCCCAACGAGAAGCGCGATCACATGGTCCTGCTGGAGTTCGTGACCGCCGCCGGGATCACTCTCGGCATGGACGAGCTGTACAAGTAA

**Tandem EPG BRET**- Vector: pcDNA 3.1 Promoter: CMV

Start-NanoLuc-EPG-EPG-mVenus-Stop

ATGGTCTTCACACTCGAAGATTTCGTTGGGGACTGGCGACAGACAGCCGGCTACAACCTGGACCAAGTCCTTGAACAGGGAGGTGTGTCCAGTTTGTTTCAGAATCTCGGGGTGTCCGTAACTCCGATCCAAAGGATTGTCCTGAGCGGTGAAAATGGGCTGAAGATCGACATCCATGTCATCATCCCGTATGAAGGTCTGAGCGGCGACCAAATGGGCCAGATCGAAAAAATTTTTAAGGTGGTGTACCCTGTGGATGATCATCACTTTAAGGTGATCCTGCACTATGGCACACTGGTAATCGACGGGGTTACGCCGAACATGATCGACTATTTCGGACGGCCGTATGAAGGCATCGCCGTGTTCGACGGCAAAAAGATCACTGTAACAGGGACCCTGTGGAACGGCAACAAAATTATCGACGAGCGCCTGATCAACCCCGACGGCTCCCTGCTGTTCCGAGTAACCATCAACGGAGTGACCGGCTGGCGGCTGTGCGAACGCATTCTGGCGATGAAGTGTGTACTTTTGGGATTCGCAGCAGTGATCGGATTCTTCGCGATCGCGGAGTCTCTTACCTGTAACACATGCTCAGTGAGTCTGATTGGAATATGTCTGAATCCCGCAACAGCGACTTGCTCCACCAACACATCCGTCTGCACCACAGGAAGAGCCAGTTTCACGGGCGTCCTCGGCTTCCTGGGCTTCAACTCCCAGGGCTGCACGGAGGGAGCTCAGTGTAATGGCACCGTGTCCGGGTCCATCCTGGGTGCGTCGTACACGGTCACTCAAACCTGCTGCAGCACAAACAACTGCAACCCCGTGACCAGCGGCGCCTCCTACGTCCAGATCTCCGTCAGCGCGGCCCTGAGCGCCGCCCTGCTGGCCTGCGTCTGGGGCCAGTCCGTCTACATGAAGTGTGTACTTTTGGGATTCGCAGCAGTGATCGGATTCTTCGCGATCGCGGAGTCTCTTACCTGTAACACATGCTCAGTGAGTCTGATTGGAATATGTCTGAATCCCGCAACAGCGACTTGCTCCACCAACACATCCGTCTGCACCACAGGAAGAGCCAGTTTCACGGGCGTCCTCGGCTTCCTGGGCTTCAACTCCCAGGGCTGCACGGAGGGAGCTCAGTGTAATGGCACCGTGTCCGGGTCCATCCTGGGTGCGTCGTACACGGTCACTCAAACCTGCTGCAGCACAAACAACTGCAACCCCGTGACCAGCGGCGCCTCCTACGTCCAGATCTCCGTCAGCGCGGCCCTGAGCGCCGCCCTGCTGGCCTGCGTCTGGGGCCAGTCCGTCTACATGGTGAGCAAGGGCGAGGAGCTGTTCACCGGGGTGGTGCCCATCCTGGTCGAGCTGGACGGCGACGTAAACGGCCACAAGTTCAGCGTGTCCGGCGAGGGCGAGGGCGATGCCACCTACGGCAAGCTGACCCTGAAGCTGATCTGCACCACCGGCAAGCTGCCCGTGCCCTGGCCCACCCTCGTGACCACCCTGGGCTACGGCCTGCAGTGCTTCGCCCGCTACCCCGACCACATGAAGCAGCACGACTTCTTCAAGTCCGCCATGCCCGAAGGCTACGTCCAGGAGCGCACCATCTTCTTCAAGGACGACGGCAACTACAAGACCCGCGCCGAGGTGAAGTTCGAGGGCGACACCCTGGTGAACCGCATCGAGCTGAAGGGCATCGACTTCAAGGAGGACGGCAACATCCTGGGGCACAAGCTGGAGTACAACTACAACAGCCACAACGTCTATATCACCGCCGACAAGCAGAAGAACGGCATCAAGGCCAACTTCAAGATCCGCCACAACATCGAGGACGGCGGCGTGCAGCTCGCCGACCACTACCAGCAGAACACCCCCATCGGCGACGGCCCCGTGCTGCTGCCCGACAACCACTACCTGAGCTACCAGTCCGCCCTGAGCAAAGACCCCAACGAGAAGCGCGATCACATGGTCCTGCTGGAGTTCGTGACCGCCGCCGGGATCACTCTCGGCATGGACGAGCTGTACAAGTAA

**EPG IRES BRET**- Vector: pcDNA 3.1 Promoter: CMV

Start- EPG-NanoLuc-IRES-EPG-mVenus-Stop

ATGAAGTGTGTACTTTTGGGATTCGCAGCAGTGATCGGATTCTTCGCGATCGCGGAGTCTCTTACCTGTAACACATGCTCAGTGAGTCTGATTGGAATATGTCTGAATCCCGCAACAGCGACTTGCTCCACCAACACATCCGTCTGCACCACAGGAAGAGCCAGTTTCACGGGCGTCCTCGGCTTCCTGGGCTTCAACTCCCAGGGCTGCACGGAGGGAGCTCAGTGTAATGGCACCGTGTCCGGGTCCATCCTGGGTGCGTCGTACACGGTCACTCAAACCTGCTGCAGCACAAACAACTGCAACCCCGTGACCAGCGGCGCCTCCTACGTCCAGATCTCCGTCAGCGCGGCCCTGAGCGCCGCCCTGCTGGCCTGCGTCTGGGGCCAGTCCGTCTACATGGTCTTCACACTCGAAGATTTCGTTGGGGACTGGCGACAGACAGCCGGCTACAACCTGGACCAAGTCCTTGAACAGGGAGGTGTGTCCAGTTTGTTTCAGAATCTCGGGGTGTCCGTAACTCCGATCCAAAGGATTGTCCTGAGCGGTGAAAATGGGCTGAAGATCGACATCCATGTCATCATCCCGTATGAAGGTCTGAGCGGCGACCAAATGGGCCAGATCGAAAAAATTTTTAAGGTGGTGTACCCTGTGGATGATCATCACTTTAAGGTGATCCTGCACTATGGCACACTGGTAATCGACGGGGTTACGCCGAACATGATCGACTATTTCGGACGGCCGTATGAAGGCATCGCCGTGTTCGACGGCAAAAAGATCACTGTAACAGGGACCCTGTGGAACGGCAACAAAATTATCGACGAGCGCCTGATCAACCCCGACGGCTCCCTGCTGTTCCGAGTAACCATCAACGGAGTGACCGGCTGGCGGCTGTGCGAACGCATTCTGGCCCCCTCTCCCTCCCCCCCCCCTAACGTTACTGGCCGAAGCCGCTTGGAATAAGGCCGGTGTGCGTTTGTCTATATGTTATTTTCCACCATATTGCCGTCTTTTGGCAATGTGAGGGCCCGGAAACCTGGCCCTGTCTTCTTGACGAGCATTCCTAGGGGTCTTTCCCCTCTCGCCAAAGGAATGCAAGGTCTGTTGAATGTCGTGAAGGAAGCAGTTCCTCTGGAAGCTTCTTGAAGACAAACAACGTCTGTAGCGACCCTTTGCAGGCAGCGGAACCCCCCACCTGGCGACAGGTGCCTCTGCGGCCAAAAGCCACGTGTATAAGATACACCTGCAAAGGCGGCACAACCCCAGTGCCACGTTGTGAGTTGGATAGTTGTGGAAAGAGTCAAATGGCTCTCCTCAAGCGTATTCAACAAGGGGCTGAAGGATGCCCAGAAGGTACCCCATTGTATGGGATCTGATCTGGGGCCTCGGTGCACATGCTTTACATGTGTTTAGTCGAGGTTAAAAAACGTCTAGGCCCCCCGAACCACGGGGACGTGGTTTTCCTTTGAAAAACACGATGATAATGGCCACAACCATGAAGTGTGTACTTTTGGGATTCGCAGCAGTGATCGGATTCTTCGCGATCGCGGAGTCTCTTACCTGTAACACATGCTCAGTGAGTCTGATTGGAATATGTCTGAATCCCGCAACAGCGACTTGCTCCACCAACACATCCGTCTGCACCACAGGAAGAGCCAGTTTCACGGGCGTCCTCGGCTTCCTGGGCTTCAACTCCCAGGGCTGCACGGAGGGAGCTCAGTGTAATGGCACCGTGTCCGGGTCCATCCTGGGTGCGTCGTACACGGTCACTCAAACCTGCTGCAGCACAAACAACTGCAACCCCGTGACCAGCGGCGCCTCCTACGTCCAGATCTCCGTCAGCGCGGCCCTGAGCGCCGCCCTGCTGGCCTGCGTCTGGGGCCAGTCCGTCTACATGGTGAGCAAGGGCGAGGAGCTGTTCACCGGGGTGGTGCCCATCCTGGTCGAGCTGGACGGCGACGTAAACGGCCACAAGTTCAGCGTGTCCGGCGAGGGCGAGGGCGATGCCACCTACGGCAAGCTGACCCTGAAGCTGATCTGCACCACCGGCAAGCTGCCCGTGCCCTGGCCCACCCTCGTGACCACCCTGGGCTACGGCCTGCAGTGCTTCGCCCGCTACCCCGACCACATGAAGCAGCACGACTTCTTCAAGTCCGCCATGCCCGAAGGCTACGTCCAGGAGCGCACCATCTTCTTCAAGGACGACGGCAACTACAAGACCCGCGCCGAGGTGAAGTTCGAGGGCGACACCCTGGTGAACCGCATCGAGCTGAAGGGCATCGACTTCAAGGAGGACGGCAACATCCTGGGGCACAAGCTGGAGTACAACTACAACAGCCACAACGTCTATATCACCGCCGACAAGCAGAAGAACGGCATCAAGGCCAACTTCAAGATCCGCCACAACATCGAGGACGGCGGCGTGCAGCTCGCCGACCACTACCAGCAGAACACCCCCATCGGCGACGGCCCCGTGCTGCTGCCCGACAACCACTACCTGAGCTACCAGTCCGCCCTGAGCAAAGACCCCAACGAGAAGCGCGATCACATGGTCCTGCTGGAGTTCGTGACCGCCGCCGGGATCACTCTCGGCATGGACGAGCTGTACAAGTAA

**EPG split NanoLuc**- Vector: pET101 Promoter: T7laq

Start-N-Terminal NanoLuc-3AA Linker-EPG-3AA Linker-C-Terminal NanoLuc-Stop

ATGGTCTTCACACTCGAAGATTTCGTTGGGGACTGGCGACAGACAGCCGGCTACAACCTGGACCAAGTCCTTGAACAGGGAGGTGTGTCCAGTTTGTTTCAGAATCTCGGGGTGTCCGTAACTCCGATCCAAAGGATTGTCCTGAGCGGTGAAAATGGGCTGAAGATCGACATCCATGTCATCATCCCGTATGAGGAGGTAGTAAGTGTGTACTTTTGGGATTCGCAGCAGTGATCGGATTCTTCGCGATCGCGGAGTCTCTTACCTGTAACACATGCTCAGTGAGTCTGATTGGAATATGTCTGAATCCCGCAACAGCGACTTGCTCCACCAACACATCCGTCTGCACCACAGGAAGAGCCAGTTTCACGGGCGTCCTCGGCTTCCTGGGCTTCAACTCCCAGGGCTGCACGGAGGGAGCTCAGTGTAATGGCACCGTGTCCGGGTCCATCCTGGGTGCGTCGTACACGGTCACTCAAACCTGCTGCAGCACAAACAACTGCAACCCCGTGACCAGCGGCGCCTCCTACGTCCAGATCTCCGTCAGCGCGGCCCTGAGCGCCGCCCTGCTGGCCTGCGTCTGGGGCCAGTCCGTCTACGGAGGTAGTGGTCTGAGCGGCGACCAAATGGGCCAGATCGAAAAAATTTTTAAGGTGGTGTACCCTGTGGATGATCATCACTTTAAGGTGATCCTGCACTATGGCACACTGGTAATCGACGGGGTTACGCCGAACATGATCGACTATTTCGGACGGCCGTATGAAGGCATCGCCGTGTTCGACGGCAAAAAGATCACTGTAACAGGGACCCTGTGGAACGGCAACAAAATTATCGACGAGCGCCTGATCAACCCCGACGGCTCCCTGCTGTTCCGAGTAACCATCAACGGAGTGACCGGCTGGCGGCTGTGCGAACGCATTCTGGCGTAA

**EPG-HaloTag**- Vector: pcDNA 3.1 Promoter: CMV

EPG Signal sequence-HaloTag-Linker-TEV site-Linker-EPG-3xFLAG

ATGAAGTGTGTACTTTTGGGATTCGCAGCAGTGATCGGATTCTTCGCGATCGCGGAGTCTATGGCAGAAATCGGTACTGGCTTTCCATTCGACCCCCATTATGTGGAAGTCCTGGGCGAGCGCATGCACTACGTCGATGTTGGTCCGCGCGATGGCACCCCTGTGCTGTTCCTGCACGGTAACCCGACCTCCTCCTACGTGTGGCGCAACATCATCCCGCATGTTGCACCGACCCATCGCTGCATTGCTCCAGACCTGATCGGTATGGGCAAATCCGACAAACCAGACCTGGGTTATTTCTTCGACGACCACGTCCGCTTCATGGATGCCTTCATCGAAGCCCTGGGTCTGGAAGAGGTCGTCCTGGTCATTCACGACTGGGGCTCCGCTCTGGGTTTCCACTGGGCCAAGCGCAATCCAGAGCGCGTCAAAGGTATTGCATTTATGGAGTTCATCCGCCCTATCCCGACCTGGGACGAATGGCCAGAATTTGCCCGCGAGACCTTCCAGGCCTTCCGCACCACCGACGTCGGCCGCAAGCTGATCATCGATCAGAACGTTTTTATCGAGGGTACGCTGCCGATGGGTGTCGTCCGCCCGCTGACTGAAGTCGAGATGGACCATTACCGCGAGCCGTTCCTGAATCCTGTTGACCGCGAGCCACTGTGGCGCTTCCCAAACGAGCTGCCAATCGCCGGTGAGCCAGCGAACATCGTCGCGCTGGTCGAAGAATACATGGACTGGCTGCACCAGTCCCCTGTCCCGAAGCTGCTGTTCTGGGGCACCCCAGGCGTTCTGATCCCACCGGCCGAAGCCGCTCGCCTGGCCAAAAGCCTGCCTAACTGCAAGGCTGTGGACATCGGCCCGGGTCTGAATCTGCTGCAAGAAGACAACCCGGACCTGATCGGCAGCGAGATCGCGCGCTGGCTGTCGACGCTCGAGATTTCCGGCGAGCCAACCACTGAGGATCTGTACTTTCAGAGCGATAACCTTACCTGTAACACATGCTCAGTGAGTCTGATTGGAATATGTCTGAATCCCGCAACAGCGACTTGCTCCACCAACACATCCGTCTGCACCACAGGAAGAGCCAGTTTCACGGGCGTCCTCGGCTTCCTGGGCTTCAACTCCCAGGGCTGCACGGAGGGAGCTCAGTGTAATGGCACCGTGTCCGGGTCCATCCTGGGTGCGTCGTACACGGTCACTCAAACCTGCTGCAGCACAAACAACTGCAACCCCGTGACCAGCGGCGCCTCCTACGTCCAGATCTCCGTCAGCGCGGCCCTGAGCGCCGCCCTGCTGGCCTGCGTCTGGGGCCAGTCCGTCTACGACTACAAGGATGACGATGACAAGGATTACAAAGACGACGATGATAAGGACTATAAGGATGATGACGACAAA

**EPG Split APEX2** Vector: pcDNA 3.1 Promoter: CMV

Start-AP Fragment-Linker-EPG-Linker-EX Fragment-Stop

ATGGGGAAATCATACCCAACAGTGTCCGCAGACTACCAGGATGCCGTGGAGAAAGCCAAGAAGAGGCTGGGAGGGTTTATCGCAGAAAAGAGGTGCGCACCTCTGATGCTGAGACTGGCTTTCCACAGCGCAGGCACCTTTGACAAGAGAACCAAAACAGGCGGACCCTTTGGAACAATCAGGTACCCCGCTGAACTGGCACATAGTGCCAACAGCGGGCTGGACATCGCCGTGCGGCTGCTGGAACCTCTGAAAGCAGAGTTCCCAATTCTGTCCTACGCCGATTTTTATCAGCTGGCAGGAGTGGTCGCTGTGGAGGTCACTGGGGGCCCCAAGGTGCCTTTTCACCCAGGACGGGAGGACAAGCCAGAACTACCTCCAGAGGGGCGCCTGCCAGATCCGACAAAGGGCTCCGACCATCTGCGAGATGTGTTTGGGAAAGCTATGGGCCTGACTGACCAGGATATCGTCGCACTGTCTGGAGGGCACACCCTTGGCGCCGCTCATAAGGAAAGGTCAGGCTTCGAGGGACCCTGGACAAGCAACCCCCTGGTTTTCGACAATTCTTACTTTACTGAACTGCTGAGTGGAGAGAAGGAAAAGGGCTCGGGCTCGACCTCGAAGTGTGTACTTTTGGGATTCGCAGCAGTGATCGGATTCTTCGCGATCGCGGAGTCTCTTACCTGTAACACATGCTCAGTGAGTCTGATTGGAATATGTCTGAATCCCGCAACAGCGACTTGCTCCACCAACACATCCGTCTGCACCACAGGAAGAGCCAGTTTCACGGGCGTCCTCGGCTTCCTGGGCTTCAACTCCCAGGGCTGCACGGAGGGAGCTCAGTGTAATGGCACCGTGTCCGGGTCCATCCTGGGTGCGTCGTACACGGTCACTCAAACCTGCTGCAGCACAAACAACTGCAACCCCGTGACCAGCGGCGCCTCCTACGTCCAGATCTCCGTCAGCGCGGCCCTGAGCGCCGCCCTGCTGGCCTGCGTCTGGGGCCAGTCCGTCTACGGATCCAAGGGCTCGGGCTCGACCTCGGGCTCGGGCGGGCTGCTGCAGCTGCCCAGCGACAAAGCCCTGCTGTCCGATCCCGTGTTCAGACCTCTGGTCGATAAGTATGCAGCCGACGAGGATGCTTTTTTCGCAGATTACGCAGAAGCACATCAGAAGCTGTCAGAACTGGGATTTGCCGACGCCTAA

**EPG Split HSVTK** Vector: pTwist CMV Puro Promoter: CMV

Start-N-Terminal HSV-TK-Linker-EPG-Linker-C-Terminal HSV-TK-Stop

ATGGCCTTGACGCCCCAGGGTGCCGAGCCCCAGAGCAACGCGGGCCCACGACCCCATATCGGGGAAACGTTATTTACCCTGTTTCGGGCCCCCGAGTTGCTGGCCCCCAACGGCGACCTGTACAACGTGTTTGCCTGGGCCTTGGACGTCTTGGCCAAACGCCTCCGTCCCATGCACGTCTTTATCCTGGATTACGACCAATCGCCCGCCGGCTGCCGGGACGCCCTGCTGCAACTTACCTCCGGGATGGTCCAGACCCATGTCACCACCCCAGGCTCCATACCGACGATCTGCGACCTGGCGCGCACGTTTGCCCGGGAGATGGGGGAGGCTCACGGAGGTGGTGGAAGTAAGTGTGTACTTTTGGGATTCGCAGCAGTGATCGGATTCTTCGCGATCGCGGAGTCTCTTACCTGTAACACATGCTCAGTGAGTCTGATTGGAATATGTCTGAATCCCGCAACAGCGACTTGCTCCACCAACACATCCGTCTGCACCACAGGAAGAGCCAGTTTCACGGGCGTCCTCGGCTTCCTGGGCTTCAACTCCCAGGGCTGCACGGAGGGAGCTCAGTGTAATGGCACCGTGTCCGGGTCCATCCTGGGTGCGTCGTACACGGTCACTCAAACCTGCTGCAGCACAAACAACTGCAACCCCGTGACCAGCGGCGCCTCCTACGTCCAGATCTCCGTCAGCGCGGCCCTGAGCGCCGCCCTGCTGGCCTGCGTCTGGGGCCAGTCCGTCTACCCCGCTCCAGCACCTGCTTCGTACCCCTGCCATCAACACGCGTCTGCGTTCGACCAGGCTGCGCGTTCTCGCGGCCATAGCAACCGACGTACGGCGTTGCGCCCTCGCCGGCAGCAAGAAGCCACGGAAGTCCGCCCGGAGCAGAAAATGCCCACGCTACTGCGGGTTTATATAGACGGTCCCCACGGGATGGGGAAAACCACCACCACGCAACTGCTGGTGGCCCTGGGTTCGCGCGACGATATCGTCTACGTACCCGAGCCGATGACTTACTGGCGGGTGCTGGGGGCTTCCGAGACAATCGCGAACATCTACACCACACAACACCGCCTCGACCAGGGTGAGATATCGGCCGGGGACGCGGCGGTGGTAATGACAAGCGCCCAGATAACAATGGGCATGCCTTATGCCGTGACCGACGCCGTTCTGGCTCCTCATATCGGGGGGGAGGCTGGGAGCTCACATGCCCCGCCCCCGGCCCTCACCATCTTCCTCGACCGCCATCCCATCGCCTTCATGCTGTGCTACCCGGCCGCGCGGTACCTTATGGGCAGCATGACCCCCCAGGCCGTGCTGGCGTTCGTGGCCCTCATCCCGCCGACCTTGCCCGGCACAAACATCGTGTTGGGGGCCCTTCCGGAGGACAGACACATCGACCGCCTGGCCAAACGCCAGCGCCCCGGTGAGCGGCTTGACCTGGCTATGCTGGCCGCGATTCGCCGCGTTTACGGGCTACTTGCCAATACGGTGCGGTATCTGCAGTGCGGCGGGTCGTGGCGGGAGGATTGGGGACAGCTTTCGGGGACGTGA

1.	Manly, B.F.J. Randomization Tests, 4th Edition by Eugene S. Edgington, Patrick Onghena.
